# Tissue dependent expression of bitter receptor *TAS2R38* mRNA

**DOI:** 10.1101/293399

**Authors:** Jennifer E. Douglas, Cailu Lin, Corrine J. Mansfield, Charles J. Arayata, Beverly J. Cowart, Andrew I. Spielman, Nithin D. Adappa, James N. Palmer, Danielle R. Reed, Noam A. Cohen

**Affiliations:** Department of Otorhinolaryngology, Head and Neck Surgery, University of Pennsylvania; Monell Chemical Senses Center; Department of Basic Science and Craniofacial Biology, New York University College of Dentistry; Philadelphia Veterans Affairs Medical Center Surgical Services

## Abstract

*TAS2R38* is a human bitter receptor gene with a common but inactive allele and people homozygous for the inactive form cannot perceive certain bitter compounds. The frequency of the inactive and active form of this receptor is nearly equal in many human populations and heterozygotes (with one copy of the active form and another copy of the inactive form) have the most common diplotype. However, heterozygotes differ markedly in their perception of bitterness, in part perhaps because of differences in *TAS2R38* mRNA expression. Other tissues including the nasal sinuses express this receptor, where it contributes to pathogen defense. We asked whether heterozygotes with high *TAS2R38* mRNA expression in taste tissue were also likely to express more *TAS2R38* mRNA in sinonasal tissue. To that end, we measured *TAS2R38* bitter taste receptor mRNA by qPCR in biopsied tissue, and learned that expression amount of one is not related to the other. We confirmed the general independence of expression in other tissue expressing *TAS2R38* mRNA using autopsy data from the GTEx project. Taste tissue *TAS2R38* mRNA expression among heterozygotes is unlikely to predict expression in other tissues, perhaps reflecting tissue-dependent function and hence regulation of this protein.

---

Human bitter taste receptors are from a gene family with 25 members that investigators originally discovered in taste receptor cells. Their protein products are the first point of contact for bitter-tasting foods and medicines, and they initiate the signal cascade that results in the perception of bitter tastes [1–3]. In addition to the taste tissue where they mediate bitter taste perception, these receptors are expressed in other tissues where they serve diverse physiological and pathophysiological roles [4]. For instance, ciliated cells in the human nasal sinuses use the bitter receptors to sense bacterial compounds, triggering an immune response [5]. One of the bitter receptors responsible for this response (T2R38, encoded by the *TAS2R38* gene) also has a well-understood role in the taste system. This gene has two main genotypes, a taster and non-taster form, that are almost equally common in most human populations, with the majority of people being heterozygous [6–8]. Heterozygous people are able to perceive the bitterness of the ligand, albeit at a reduced intensity relative to people who are homozygous for the taster form [9]. However, heterozygous people vary considerably in their sensitivity to bitter ligands, which could be due in part to variable mRNA expression of the active form of the receptor [10]. This point may have practical importance because clinicians are exploring the use of taste testing as means of understanding sinonasal disease risk and surgical outcomes [11]. It would be helpful to know if heterozygous patients who experience more taste bitterness are likely to express more *TAS2R38* mRNA in nasal tissue. Therefore, we determined whether the range of mRNA expression in the taste cells is similar to that expressed in the sinonasal cells, focusing on heterozygotes. To that end, we biopsied taste and sinonasal tissue from genetically characterized adults and measured mRNA expression; in addition, each person was rated the bitterness rating of a T2R38 ligand and other taste solutions. To understand whether these results generalize to other tissues, we similarly analyzed gene expression data available from the GTEx consortium [12].

## Methods

### Subjects

Most of our subjects were patients of one of the authors (NAC), an attending surgeon in the Philadelphia Veterans Affairs Medical Center and the Department of Otorhinolaryngology – Head and Neck Surgery at the University of Pennsylvania. They were undergoing sinonasal surgery for reasons other than chronic rhinosinusitis (e.g., reconstructive nasal surgery, tear duct or sinus surgery to access tumors in the cranial vault/skull base). We recruited additional subjects from within those departments who did not undergo surgery, likewise excluding those with a history of chronic rhinosinusitis. We did not exclude smokers because of the high rates among veterans, e.g., [13].

### Sample processing

For genotyping, we collected saliva using Oragene-Discover OGR-500 collection kits (DNA Genotek, Ottawa, Ontario, Canada), and we extracted genomic DNA following manufacturer directions. We genotyped alleles of the *TAS2R38* gene (accession number NM_176817) using real-time PCR single-nucleotide polymorphism genotyping assays for rs713598, rs1726866, and rs10246939 (C___8876467_10, C___9506827_10, C___9506826_10, respectively; TaqMan^TM^ from Thermo Fisher, Waltham, MA) with the QuantStudio 12K Flex or StepOnePlus sequence detection system (Applied Biosystems, Grand Island, NY). We assigned *TAS2R38* diplotypes (the combination of haplotypes) for each person as previously described [9]. See Table 1 for haplotype details.

**Table 1.** Common variants of *TAS2R38*.

| TAS2R38 | A49P | V262A | I296V | Haplotype |
| --- | --- | --- | --- | --- |
| Taster | PP | AA | VV | PAV |
| Non-taster | AA | VV | II | AVI |

For mRNA analysis we collected taste samples (fungiform papillae) following published procedures [14], and we collected nasal tissue either by brush (MasterAmp Buccal Brush, Epicentre, Madison, WI) or by biopsy of the inferior turbinate (Figure 1). We noted the time of tissue collection and placed the tissue samples in RNA preservative (RNAlater, Invitrogen, Life Technologies, Carlsbad, CA). To isolate the RNA from all tissue samples (average of six taste papillae and 31 mg nasal tissue per subject), we followed the manufacturer’s directions; for processing the taste tissue and nasal brush samples, we used Quick-RNA MiniPrep R1054, Zymo Research; for the nasal biopsy samples, we used ZR-Duet kit. We evaluated RNA quality expressed as an RNA integrity number (RIN) using the Agilent 2200 TapeStation system (Agilent Technologies, Santa Clara, CA). We synthesized cDNA using the Ovation RNA Amplification System V2 (cat. no. 3100-12, NuGEN Technologies, San Carlos, CA), including a DNase step (Zymo Research). We purified the resulting cDNA using QIAquick PCR Purification Kit (cat. no. 28104, Qiagen, Germantown, MD).

**Figure 1.**
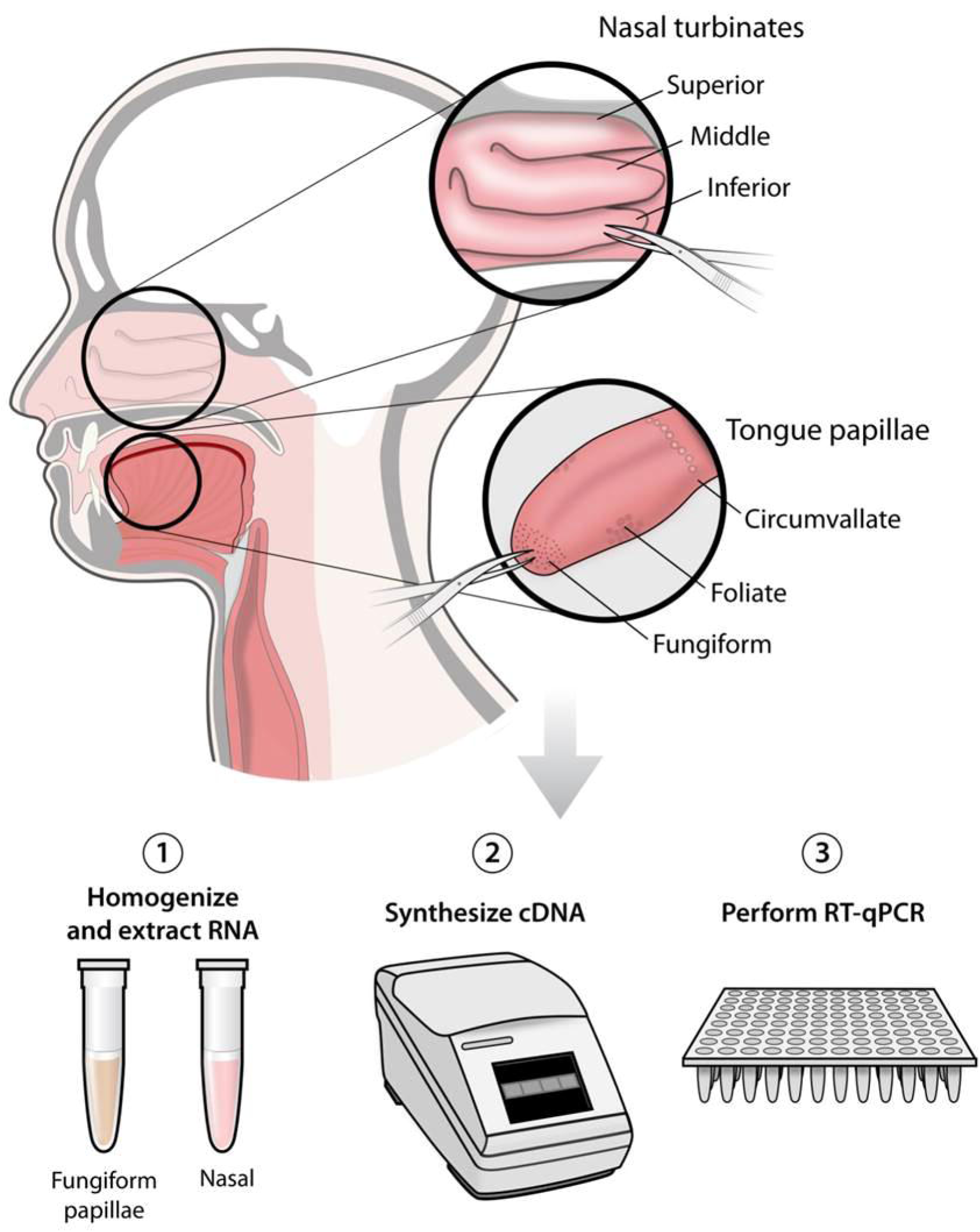
Schematic of the tissue collection, processing and qPCR analysis.

### Cell-specific assays and allele-specific TAS2R38 cDNA measures

We used allele-specific probes for the*TAS2R38* variant site from the assays listed above. We also measured the cell-specific markers *CFTR* (Hs00357011_m1) for motile cilia in nasal tissue and *GNAT3* (Hs01385403_m1) for type 2 taste cells in taste tissue, as well as *GAPDH* (cat. no. 4326317E) as a housekeeping gene in both tissues. (Bitter receptors are expressed in a subset of type 2 taste cells). The bitter taste receptor genes have a single exon, so we could not use intron-spanning primers; therefore, we verified absence of genomic DNA in the cDNA samples by size-specific primers from the *ABL1* gene [15]. We excluded cDNA samples with consistent evidence of residual genomic DNA from some analyses as explained below.

### RNA-seq

*TAS2R38* mRNA is expressed in only a small fraction of the cells within the biopsies, and measurement of this low abundance transcript can be imprecise. Therefore, on samples that met the quality standards of the Next-Generation Sequencing Core of the University of Pennsylvania (RIN > 7), their staff performed library preparation and sequencing (100 bp single-end) on the HiSeq 4000 sequencer (Illumina, San Diego, CA) using the manufacturer’s sequencing protocols. We mapped reads to the reference genome (GRCh38.p10) after the raw sequence data in fastq format passed the quality filters of *Trimmomatic* [16] and we normalized the counts using the R package *ballgown* [17]. We compared the normalized counts to the expression measured in qPCR and we examined the abundance of associated taste signaling genes as a general validation step.

### Taste testing and dietary measures

Using published methods [11, 18] and prior to biopsy, we asked subjects to rate the intensity of several taste solutions, including phenylthiocarbamide (PTC), a ligand of the T2R38 receptor [4]. Each subject rated six taste solutions twice: water, PTC (180 μM), denatonium benzoate (1.8 μM), quinine (56 μM), sodium chloride (0.25 M) and sucrose (0.35 M). People also chose a quality descriptor for each solution from among the following list: salty, sour, bitter, sweet, and no flavor. (Detailed taste methods are available online at https://osf.io/hn87s/). We also asked subjects to quantify their habitual caffeine consumption using a modified version of the Harvard Food Frequency Questionnaire, a semi-quantitative dietary assessment tool [19, 20] (available online at https://osf.io/j3suy/). We included questions about caffeine consumption because prior studies suggested that subjects who consumed more caffeine have higher expression of bitter taste receptors in taste papillae [10, 20].

### Analysis

The abundance of each *TAS2R38* allele was expressed relative to the housekeeping gene [21] using the median of three reporters as input values, with the three variant sites averaged as previously described [10]. If we detected no expression, we used a cycle threshold value of 40. We excluded from some analyses samples that did not contain the target cell type and we evaluated the correlation of gene expression in taste vs nasal tissue using a Spearman correlation coefficient; we also used this statistical method to compute the correlation between qPCR and RNA-seq values. For all analyses listed above and below, we computed the statistical tests with R (version 3.4.2) and R-studio (version 1.1.383).

### Ethics and reproducibility

The institutional review boards of Philadelphia Veterans Affairs Medical Center and the University of Pennsylvania approved these protocols. In advance of data collection, we registered our hypothesis at the Center for Open Science (Open Science Framework, https://osf.io/xbp6q/). All data and analysis scripts are available for download at Github (https://github.com/DanielleReed/R21) and the Center for Open Science (https://osf.io/yeqjf).

### GTEx data

After we obtained appropriate approvals, administrators from the GTEx project provided RNA-seq data from 738 people from post-mortem tissue samples (51 tissues; GTEx; #12732: Bitter receptor gene expression: patterns across tissues). We extracted the normalized RPKM (reads per kilobase of transcript) values for the *TAS2R38* gene across the tissues. We classified samples as expressing *TAS2R38* mRNA if the RPKM was above 0 and computed the percentage of samples that expressed TAS2R8 mRNA by tissue. For samples where this gene was expressed, we tested whether it was affected by *TAS2R38* genotype with a Kruskal-Wallis test. We examined whether people expressing higher amounts in one tissue were likely to expression higher amounts in other tissues with Spearman correlational analysis.

## Results

### Screening and subjects

Forty-five adult subjects provided samples for *TAS2R38* genotyping during a preoperative or other clinic visit, from which we identified a sample enriched for heterozygotes (AVI/PAV; N=19) to provide nasal epithelial tissue and taste tissue from the fungiform papillae. We also recruited four PAV/PAV and four AVI/AVI subjects for comparison. In all, we obtained tissue and performed sensory testing on 27 subjects: 23 males and 4 females. Sixteen subjects (59%) were of European descent, and the remaining were racially diverse; which reflects the metropolitan area from which the sample was collected (Philadelphia, PA) [22]. We list individual subjects and their characteristics in Supplemental Table 1. The median age of the subjects was 35 years (range, 23-71 years). We depicted the subjects and procedures in Figure 2.

**Figure 2.**
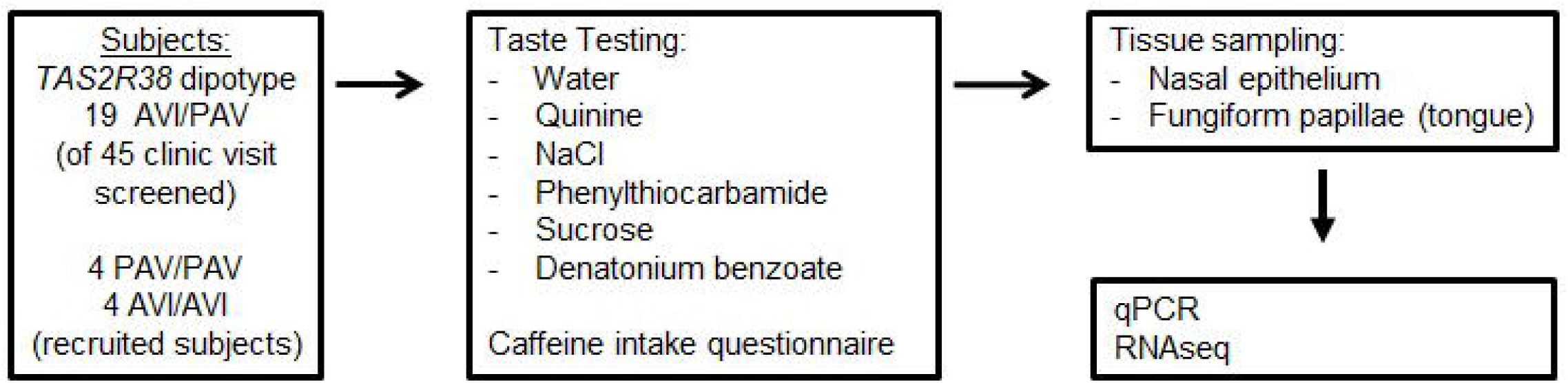
Experimental flowchart.

All subjects provided fungiform papillae by biopsy and all subjects provided nasal tissue, 15 by nasal brush and 12 by biopsy. The RIN for the taste samples were noticeably higher than for the nasal samples [t(26)=5.06, p=2.857 x 10^-5^], with the median RIN of 6.4 for the taste samples and 2.2 for nasal samples overall, with an advantage for the biopsy (RIN=2.6) over the brush method (RIN=1.4; Supplemental Table 2). However, people providing taste samples that produced the highest quality RNA were not more likely to provide high-quality nasal samples (Spearman r=0.10, p=0.78; Supplemental Figure 1). Despite the difference in RNA quality between taste and nasal tissues, observed C_t_(cycle threshold) for the housekeeping gene *GAPDH* did not differ between these two tissues [t(26)=1.10, p=0.2794]. (Cycle threshold ranges from 0 to 40 and is an indirect measure of gene abundance).

Of the 27 taste samples collected, 13 had detectable amounts of the cell-specific marker *GNAT3*; of the nasal samples, 19 had detectable amounts of the cell-specific marker *CFTR*. Nine subjects had samples with appropriate cell-specific markers in both tissues. Eight of these nine samples were from people who are *TAS2R38* heterozygotes, but we excluded two additional samples because we consistently detected residual genomic DNA ( Supplemental Table 2). Among these six people, there was no relationship between *TAS2R38* PAV mRNA abundance from taste versus nasal tissue (r=0.2, p=0.70, Spearman; Figure 3A). The pattern of results was similar when we included all 19 heterozygous subjects in the analysis (Supplemental Figure 3A). Our pre-registered hypothesis which was that heterozygotes with higher *TAS2R38* mRNA in taste tissue would also have higher expression in nasal tissue was not confirmed.

**Figure 3.**
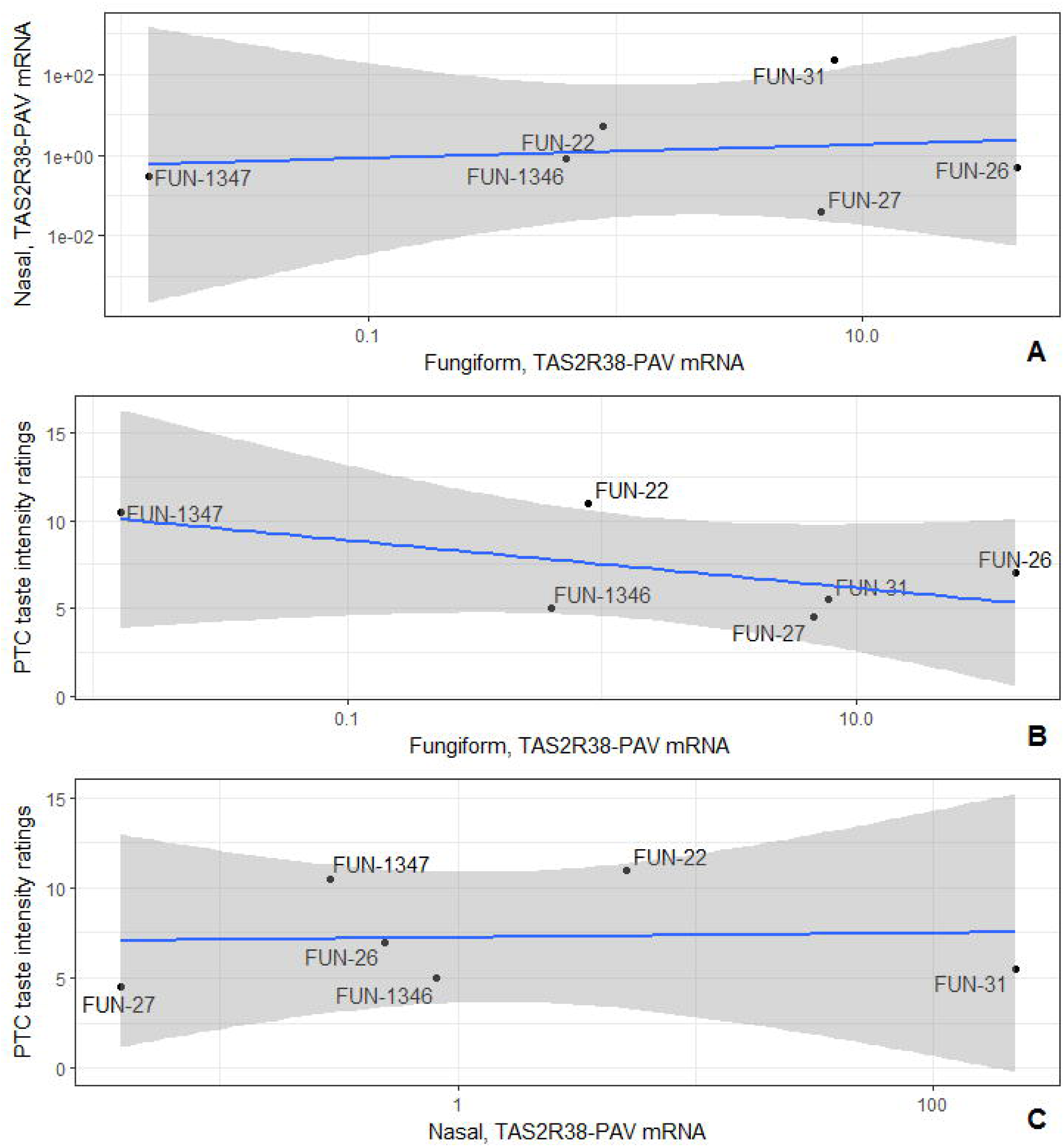
Correlation among abundance of the *TAS2R38* PAV mRNA in nasal, taste tissue, and PTC taste intensity. Each dot represents data from one subject (subject ID prefix “FUN”). The blue line shows linear correlation; the gray shading shows the confidence interval. Axes scales are the log of the relative abundance of the *TAS2R8* PAV mRNA relative to *GAPDH*. Six heterozygous people provided samples in which we could detect appropriate cell-specific markers and no residual genomic DNA. Abundance of the *TAS2R38* PAV mRNA in nasal versus taste tissue (A) and PTC taste intensity ratings vs abundance of the *TAS2R38* PAV mRNA in taste tissue (B) or nasal (C).

For RNAseq analysis, six samples, all of which were from taste and none from nasal tissue, were of sufficient quality for sequencing (RIN>7; Supplemental Table 2). While there were too few samples to draw meaningful statistical comparisons, we noted that the expression of *TAS2R38* mRNA is similar between the two methods commonly used to measure gene expression, RNAseq and qPCR (r=0.71, p=0.11, Spearman; Supplement Figure 2). As expected, we also detected gene expression for markers of taste receptor cells, *TRPM5* and *GNAT3*, albeit at a low abundance ( Supplemental Table 3).

People with one or more PAV haplotypes rated PTC as more intense than did those homozygous for the inactive form [F(2, 24)=7.13, p=0.0037; Supplemental Figure 4]. There was no relationship between *TAS2R38* PAV mRNA abundance from fungiform or nasal tissue and ratings of PTC bitterness intensity (fungiform, r=-0.20, p=0.70; nasal, r=0.31, p=0.54; Spearman; Figure 3B, C). For the 19 heterozygous subjects (which includes those with no cell-specific expression), we observed a similar result (fungiform, r=-0.43, p=0.06; nasal, r=-0.04, p=0.88; Spearman; Supplemental Figure 3B, C). The negative correlation between taste tissue mRNA and sensory measures (−0.43) is in the opposite direction from that which we observed in an earlier published study [10].

**Figure 4.**
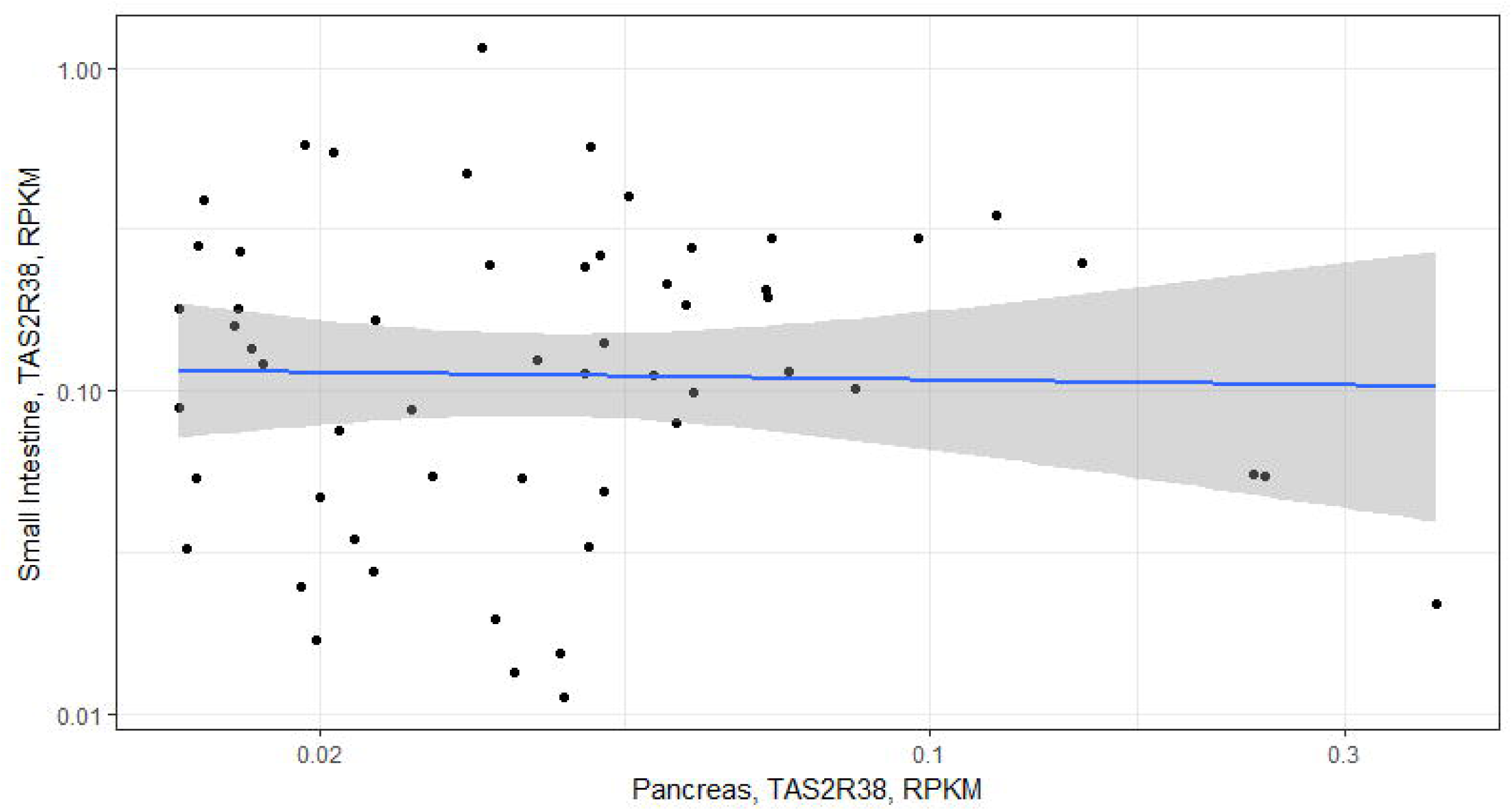
A scatterplot for the correlation analysis of the *TAS2R38* mRNA expression between small intestine and pancreas from the GTEx database.

The amount of caffeine drunk by the six heterozygous subjects that met the most stringent inclusion criterion (Figure 3A; Supplemental Table 2) was also unrelated to *TAS2R38* PAV mRNA abundance (r=0.37, p=0.47, Spearman). Other taste ratings were as expected, but with a few anomalies ( Supplemental Table 4). All subjects described sucrose as sweet, and all but one subject described sodium chloride as salty. A number of subjects reported the bitter stimuli to be sour, but this is a common error [23]. However, a few people rated the water as having a flavor ( Supplemental Table 5). This incorrect rating was for the second but not first trial of water testing, and may be due to carry-over effects from preceding taste stimuli.

Subjects differed in fungiform papillae density, with values ranging from 0 to 6 papillae per cubic centimeter, but those with a higher density did not rate PTC as more bitter than did those with a lower density (r=-0.05, p=0.81; Spearman). For the RNAseq data, the mRNA expression of *TAS2R38* in taste was not correlated with that of *TRPM5* (r=-0.03, p=0.96; Spearman) or *GNAT3* (r=0.029, p=1.0; Spearman).

Drawing on the GTEx dataset, we learned that *TAS2R38* mRNA could be detected in a low of 5% of the samples for some tissues (e.g., muscle) and as high as 77% in the small intestine, although not all tissues had complete data and thus were less informative than others ( Supplemental Table 6). For samples that expressed detectable amounts of *TAS2R38* mRNA, there was a >300-fold range (RPKM values of 0.0036 vs 1.159), with the highest RPKM values in small intestine, testis, pancreas, transverse colon and bladder (Supplemental Figure 6). For these tissues, people did not differ in the *TAS2R38* mRNA expression by diplotype (Kruskal-Wallis, p>0.05; Supplemental Figure 6) so all subjects were included in the remaining analysis. Using the tissues with higher *TAS2R38* mRNA expression, we learned that the abundance in expression in one tissue was unrelated to abundance in the other tissues (e.g., small intestine vs pancreas: r=0.08, p=0.57; Figure 4, Supplemental Figure 7) except for the small intestine and colon (Supplemental Figure 8). We omitted one tissue, bladder, from this correlational analysis because there were too few tissue-pairs for meaningful comparisons, e.g., N=2-3.

## Discussion

The *TAS2R38* gene codes for the bitter taste receptor protein T2R38 and has two common forms that predict whether people can taste the ligand PTC at low concentration. However, the majority of people are heterozygotes, and their ability to taste PTC ranges widely. This observation is important medically because PTC sensitivity may predict disease susceptibility and surgical outcome for sinonasal disease [5, 11, 24, 25] and potentially predicts other conditions such as dental caries [26–30]. Previously, we demonstrated that heterozygous people rated the bitterness of a T2R38 ligand as more intense if they had higher expression of the taster allele [10]. We were motivated to conduct the current research in part by the practical usefulness of knowing whether taste ratings by heterozygotes would predict the abundance of *TAS2R38* mRNA in the nasal sinuses and a more vigorous immune response. This information would in turn help us to understand their disease risk and surgical outcomes. However, we learned that it did not—higher gene expression in taste tissue did not predict higher expression in sinonasal tissue. We also learned that this observation appears to be a general principle for other tissues in the body that express *TAS2R38* mRNA, which is broadly consistent with the pattern of gene expression for many other genes, where abundance depends on the particular tissue more so than the particular individual from which it comes [12]. Taste testing remains a highly effective way to estimate *TAS2R38* genotype, but it does not predict *TAS2R38* mRNA gene expression in the nasal sinuses, at least among the people studied here.

This study has limitations that arise from RNA quality and the low expression of gene generally. We found marked differences between the nasal and taste tissue in RNA quality. RNA quality was low in a *few* taste samples but it was low or very low in nearly *all* nasal samples. Other investigators have observed that some tissues have consistently lower RNA quality than others do [31], but it appears that nasal tissue is at the low extreme, possibly due to the presence of RNA-degrading enzymes within the nasal mucus [32, 33]. Furthermore, only a few cells within the human taste [34] and nasal [5] tissue express the bitter taste receptor gene and therefore abundance in the tissue overall is very low, near the level of detection for qPCR. However, the results from RNAseq (a method which can detect a greater range of expression than qPCR) partially offset this limitation and the results from these two methods agreed, although we were only able to obtain RNAseq data from a subset of samples owing to low RNA quality.

To determine if this independence of *TAS2R38* mRNA in taste and nasal tissues followed a general rule or was an exception, we also examined data from the GTEx consortium. Consortium investigators collected and measured RNA expression in tissues from many people after death. While *TAS2R38* mRNA expression is undetectable in most tissues, we focused on tissues with higher expression, e.g., small intestine and pancreas. In the small intestine, extrapolating from observations in the mouse, ligands of T2R38 initiate hormone release from enteroendocrine cells that slows nutrient absorption [35]. We know little about pancreatic *TAS2R38* mRNA expression or its T2R38 function except that, according to the Human Protein Atlas, the protein is present in exocrine glandular cells [36]. It appears that tissues that express *TAS2R38* mRNA do so independently, and higher expression in one tissue does not predict higher expression in a different tissue, similar to our observation in taste and nasal tissue. We also note a mismatch between low *TAS2R38* mRNA patterns in the GTEx dataset, with the more abundant protein abundance in the Human Protein Atlas dataset. This mismatch between mRNA and protein abundance (albeit from two different large-scale data repositories) point to translation of mRNA to protein as an uncharacterized but potentially important regulatory step.

In contrast to our earlier research outcomes, we did not observe a relationship between ratings of PTC bitterness intensity and *TAS2R38* mRNA [10], nor did we observe a relationship between gene expression and caffeine consumption [20]. Likewise, we also found no correlation between taste papillae density and bitterness perception [37, 38]. We have no explanation for these discrepancies except that sensory testing of clinic patients may differ from testing in experimental psychology laboratories, e.g., clinic testing engenders more anxiety and less attention to the testing procedures or differences in the sensory rating scales. The subjects themselves were markedly different too, healthy young subjects in the first study and older patients studied here. However, the relationship between *TAS2R38* genotype and PTC bitterness was clear, and consistent with many other sensory studies of T2R38 ligands [9, 39–42]. Even with these caveats, the results show clearly that deficiencies in the sensory testing, if present, were not large.

## Acknowledgements

Author responsibilities were as follows: D.R.R., N.A.C., and J.E.D. designed the research; C.L. and D.R.R. analyzed data, performed statistical analysis, and wrote the manuscript; J.E.D. analyzed data and wrote the manuscript; C.J.A. analyzed data and performed statistical analysis; J.E.D. and C.J.M. conducted the research, A.I.S. and B.J.C. consulted on the research and manuscript; N.A.C. and D.R.R. share primary responsibility for the final content. All authors read and approved the final manuscript.

## Funding

The National Institutes of Health (NIH) supported this research with award NIH R21DC013886 (NAC and DRR). The National Institute on Deafness and Other Communication Disorders Administrative Research Supplement to Promote Emergence of Clinician-Scientists in Chemosensory Research provided support for Jennifer E. Douglas. We collected genotype data from equipment purchased in part with NIH funds from OD018125.

---

**Supplemental Table 1:** Subject characteristics

| Subject Id | Sex | Race | TAS2R38 Diplotype |
| --- | --- | --- | --- |
| FUN-10 | M | 1 | PAV/PAV |
| FUN-12 | M | 2 | AVI/PAV |
| FUN-13 | F | 1 | AVI/PAV |
| FUN-1334 | M | 9 | PAV/PAV |
| FUN-1335 | M | 9 | AVI/PAV |
| FUN-1336 | M | 4 | AVI/AVI |
| FUN-1337 | M | 1 | AVI/AVI |
| FUN-1340 | M | 1 | AVI/PAV |
| FUN-1341 | M | 1 | PAV/PAV |
| FUN-1342 | M | 4 | AVI/PAV |
| FUN-1343 | M | 1 | AVI/PAV |
| FUN-1344 | M | 9 | AVI/AVI |
| FUN-1346 | M | 9 | AVI/PAV |
| FUN-1347 | M | 1 | AVI/PAV |
| FUN-1349 | M | 4 | AVI/PAV |
| FUN-16 | M | 1 | AVI/PAV |
| FUN-17 | F | 1 | AVI/AVI |
| FUN-18 | M | 6 | AVI/PAV |
| FUN-21 | M | 1 | AVI/PAV |
| FUN-22 | F | 6 | AVI/PAV |
| FUN-26 | M | 1 | AVI/PAV |
| FUN-27 | M | 1 | AVI/PAV |
| FUN-3 | M | 1 | PAV/PAV |
| FUN-31 | M | 1 | AVI/PAV |
| FUN-34 | F | 1 | AVI/PAV |
| FUN-4 | M | 1 | AVI/PAV |
| FUN-9 | M | 2 | AVI/PAV |

FUN=study code, abbreviated for fungiform taste papillae.
1 = Caucasian, 2=Asian, 4=African American, 6=Other/mixed, 9=Race not available/provided.
PAV/PAV is the active or taster form; AVI/AVI is the inactive or non-taster form.

**Supplemental Table 2:** Sample quality, inclusion and RNA integrity for fungiform and nasal samples

| Subject ID | Key | FP RIN | Nasal Tissue |  |
| --- | --- | --- | --- | --- |
|  |  |  | RIN | Sample Type |
| FUN-3 | c | <1.0 | <1.0 | Brush |
| FUN-34 | c | 7.2 | <1.0 | Brush |
| FUN-1334 | b | <1.0 | 1.0 | Biopsy |
| FUN-1335 | b | 4.2 | 1.0 | Biopsy |
| FUN-1336 | b | 4.7 | 1.0 | Biopsy |
| FUN-12 | b | <1.0 | 1.0 | Brush |
| FUN-18 | b | 7.3 | 1.0 | Brush |
| FUN-26 | a, d | 8.2 | 1.0 | Brush |
| FUN-4 | c | 4.0 | 1.0 | Brush |
| FUN-10 | c | 5.7 | 1.3 | Brush |
| FUN-1341 | a, e | 8.3 | 1.4 | Biopsy |
| FUN-17 | b, e | 8.0 | 1.4 | Brush |
| FUN-9 | b, e | 7.5 | 1.6 | Brush |
| FUN-16 | b | 8.5 | 2.2 | Brush |
| FUN-13 | a | 8.1 | 2.3 | Brush |
| FUN-1346 | a, d | 7.4 | 2.4 | Biopsy |
| FUN-1343 | b | 6.4 | 2.6 | Biopsy |
| FUN-1344 | b, e | 8.6 | 2.6 | Biopsy |
| FUN-1342 | b, e | 7.9 | 2.9 | Biopsy |
| FUN-22 | a, d, e | 7.5 | 3.0 | Brush |
| FUN-1349 | b | <1.0 | 3.2 | Biopsy |
| FUN-1347 | a, d | 8.7 | 3.3 | Biopsy |
| FUN-21 | a | 5.7 | 3.3 | Brush |
| FUN-1337 | b | 5.3 | 3.4 | Biopsy |
| FUN-31 | a, d | 2.3 | 3.7 | Brush |
| FUN-1340 | b | 6.1 | 3.8 | Biopsy |
| FUN-27 | a, d | <1.0 | 5.9 | Brush |

Table is sorted by ascending RIN of the nasal samples. FP=Fungiform papillae. RIN=RNA integrity number. Key: (a) Samples with *GNAT3* mRNA in taste tissue and *CFTR* mRNA in nasal tissue, N=9; (b) Samples with either *GNAT3* mRNA in taste tissue or *CFTR* mRNA in nasal samples but not both, N=14; (c) Subjects that provided samples that neither expressed *GNAT3* mRNA in taste tissue nor *CFTR* mRNA in nasal tissue, N = 4; (d) Subjects from group (a) that were also heterozygous (AVI/PAV) and also had no consistent evidence of genomic DNA in the sample, N=6; (e) Samples used for the RNAseq study, N=6.

**Supplemental Table 3:** RNAseq indicates biopsied tissue likely to contain taste receptor cells

| **Gene Name** | **Gene Id** | **FUN-1341** | **FUN-1342** | **FUN-1344** | **FUN-17** | **FUN-22** | **FUN-9** |
| --- | --- | --- | --- | --- | --- | --- | --- |
| *GNAT3* | ENSG00000214415 | 1.27 | 0.73 | 2.93 | 1.87 | 1.06 | 1.91 |
| *TRPM5* | ENSG00000070985 | 0.12 | 0.02 | 1.23 | 0.05 | 0.12 | 0.25 |
| *TAS2R38* | ENSG00000257138 | 0.18 | 0.78 | 1.95 | 0.106 | 0.03 | 0.00 |

TRPM5 is commonly expressed in either taste receptor cells and the protein product is part of the signaling cascade.

**Supplemental Table 4:** Mean ratings of taste solution intensity by *TAS2R38* diplotype

| Solution | Diplotype |  |  | F(2,24) | P-Value |
| --- | --- | --- | --- | --- | --- |
|  | AVI/AVI | AVI/PAV | PAV/PAV |  |  |
| Phenylthiocarbamide | 1.1 | 7.1 | 8.1 | 7.1 | 0.004** |
| Quinine | 4.8 | 2.5 | 0.9 | 3.9 | 0.034* |
| Denatonium benzoate | 11.8 | 8.6 | 7.4 | 3.0 | 0.071 |
| NaCl | 9.1 | 7.1 | 6.5 | 1.8 | 0.326 |
| Sucrose | 7.0 | 7.1 | 7.0 | 0.0 | 0.996 |
| Water | 2.0 | 0.4 | 0.0 | 9.4 | <0.001** |

\*P<0.05; \*\*p<0.001.

**Supplemental Table 5:** Individual subject ratings of solution flavors.

| Diploype | Identifier | Wat 1 | Wat 2 | Quin 1 | Quin 2 | NaCl 1 | NaCl 2 | PTC 1 | PTC 2 | Suc 1 | Suc 2 | DB 1 | DB 2 |
| --- | --- | --- | --- | --- | --- | --- | --- | --- | --- | --- | --- | --- | --- |
| AVI/AVI | FUN-1336 | No | **Bitter** | Bitter | **Salty** | Salty | Salty | No | No | Sweet | Sweet | Bitter | Bitter |
| AVI/AVI | FUN-1337 | No | **Bitter** | No | Bitter | Salty | Salty | Bitter | No | Sweet | Sweet | Bitter | Bitter |
| AVI/AVI | FUN-1344 | **Sweet** | **Bitter** | Sour | Bitter | Salty | Salty | Sweet | No | Sweet | Sweet | Bitter | Bitter |
| AVI/AVI | FUN-17 | No | No | Bitter | Bitter | Salty | Salty | No | No | Sweet | Sweet | Bitter | Bitter |
| AVI/PAV | FUN-12 | No | No | Bitter | Bitter | Salty | Salty | Bitter | Bitter | Sweet | Sweet | Bitter | Bitter |
| AVI/PAV | FUN-13 | No | No | Bitter | Bitter | Salty | Salty | Bitter | Bitter | Sweet | Sweet | Bitter | Bitter |
| AVI/PAV | FUN-1335 | No | No | Bitter | No | Salty | Salty | Bitter | Bitter | Sweet | Sweet | Bitter | Bitter |
| AVI/PAV | FUN-1340 | No | No | Salty | Bitter | Salty | Salty | Sour | Salty | Sweet | Sweet | Bitter | Sour |
| AVI/PAV | FUN-1342 | No | No | No | No | Salty | Salty | Bitter | Bitter | Sweet | Sweet | Bitter | Bitter |
| AVI/PAV | FUN-1343 | No | No | Bitter | No | **Bitter** | Salty | Bitter | Bitter | Sweet | Sweet | Sour | Sour |
| AVI/PAV | FUN-1346 | No | No | Bitter | No | Salty | Salty | Bitter | Bitter | Sweet | Sweet | Bitter | Bitter |
| AVI/PAV | FUN-1347 | **Bitter** | No | Bitter | Bitter | Salty | Salty | Bitter | Bitter | Sweet | Sweet | Bitter | Bitter |
| AVI/PAV | FUN-1349 | No | **Bitter** | No | No | Salty | Salty | Bitter | No | Sweet | Sweet | Bitter | No |
| AVI/PAV | FUN-16 | Salty | No | Bitter | No | Salty | Salty | Bitter | Bitter | Sweet | Sweet | Bitter | Sour |
| AVI/PAV | FUN-18 | No | **Bitter** | Bitter | Bitter | Salty | Salty | Bitter | Bitter | Sweet | Sweet | Bitter | Bitter |
| AVI/PAV | FUN-21 | No | No | Bitter | No | Salty | Salty | Bitter | Bitter | Sweet | Sweet | Bitter | Sour |
| AVI/PAV | FUN-22 | No | No | Bitter | Bitter | Salty | Salty | Bitter | Bitter | Sweet | Sweet | Bitter | Bitter |
| AVI/PAV | FUN-26 | No | **Bitter** | Bitter | Bitter | Salty | Salty | Bitter | Bitter | Sweet | Sweet | Bitter | Bitter |
| AVI/PAV | FUN-27 | No | No | Bitter | No | Salty | Salty | Bitter | Bitter | Sweet | Sweet | Bitter | Bitter |
| AVI/PAV | FUN-31 | No | No | Bitter | Sour | Salty | Sour | Bitter | Sour | Sweet | Sweet | Bitter | Bitter |
| AVI/PAV | FUN-34 | **Salty** | No | Bitter | Bitter | Salty | Salty | Bitter | Bitter | Sweet | Sweet | Bitter | Bitter |
| AVI/PAV | FUN-4 | No | No | No | Bitter | Salty | Salty | Bitter | Bitter | Sweet | Sweet | Bitter | Bitter |
| AVI/PAV | FUN-9 | No | No | Bitter | Bitter | Salty | Salty | Bitter | Bitter | Sweet | Sweet | Bitter | Bitter |
| PAV/PAV | FUN-10 | No | No | Bitter | No | Salty | Salty | Bitter | Bitter | Sweet | Sweet | Bitter | Bitter |
| PAV/PAV | FUN-1334 | No | No | No | No | Salty | Salty | Bitter | Sour | Sweet | Sweet | Bitter | Bitter |
| PAV/PAV | FUN-1341 | No | No | No | No | Salty | Salty | Bitter | Bitter | Sweet | Sweet | Sour | Bitter |
| PAV/PAV | FUN-3 | No | Salty | Bitter | Sour | Salty | Salty | Bitter | Bitter | Sweet | Sweet | Bitter | Bitter |

DB=denatonium benzoate. Font face indicates **incorrect** responses. Each person sampled each solution twice.

**Supplemental Table 6:** Detection of *TAS2R38* mRNA expression by tissue

| *Tissue* | *Yes* | *No* | *Per* |
| --- | --- | --- | --- |
| Small Intestine - Terminal Ileum | 111 | 32 | 77.6 |
| Testis | 210 | 70 | 75.0 |
| Pancreas | 197 | 66 | 74.9 |
| Bladder | 6 | 5 | 54.5 |
| Colon - Transverse | 148 | 137 | 51.9 |
| Whole Blood | 167 | 276 | 37.7 |
| Spleen | 62 | 107 | 36.7 |
| Fallopian Tube | 2 | 5 | 28.6 |
| Minor Salivary Gland | 25 | 78 | 24.3 |
| Lung | 94 | 349 | 21.2 |
| Cervix - Endocervix | 1 | 4 | 20.0 |
| Kidney - Cortex | 10 | 40 | 20.0 |
| Skin - Sun Exposed (Lower leg) | 91 | 417 | 17.9 |
| Cervix - Ectocervix | 1 | 5 | 16.7 |
| Prostate | 25 | 131 | 16.0 |
| Skin - Not Sun Exposed (Suprapubic) | 60 | 346 | 14.8 |
| Artery - Coronary | 23 | 159 | 12.6 |
| Thyroid | 58 | 418 | 12.2 |
| Colon - Sigmoid | 30 | 223 | 11.9 |
| Brain - Cortex | 20 | 154 | 11.5 |
| Pituitary | 21 | 170 | 11.0 |
| Stomach | 29 | 240 | 10.8 |
| Vagina | 11 | 113 | 8.9 |
| Brain - Anterior cingulate cortex (EB24) | 11 | 117 | 8.6 |
| Uterus | 10 | 106 | 8.6 |
| Adipose - Visceral (Omentum) | 31 | 332 | 8.5 |
| Brain - Cerebellum | 16 | 176 | 8.3 |
| Brain - Spinal cord (cervical c-1) | 8 | 89 | 8.2 |
| Brain - Nucleus accumbens (basal ganglia) | 13 | 147 | 8.1 |
| Nerve - Tibial | 35 | 405 | 8.0 |
| Ovary | 11 | 127 | 8.0 |
| Brain - Frontal Cortex (EB9) | 11 | 132 | 7.7 |
| Esophagus - Mucosa | 32 | 406 | 7.3 |
| Breast - Mammary Tissue | 22 | 282 | 7.2 |
| Cells - Transformed fibroblasts | 26 | 334 | 7.2 |
| Liver | 13 | 172 | 7.0 |
| Artery - Tibial | 32 | 430 | 6.9 |
| Brain - Hippocampus | 9 | 121 | 6.9 |
| Brain - Putamen (basal ganglia) | 9 | 126 | 6.7 |
| Brain - Caudate (basal ganglia) | 11 | 163 | 6.3 |
| Esophagus - Muscularis | 23 | 366 | 5.9 |
| Cells - EEV-transformed lymphocytes | 8 | 130 | 5.8 |
| Heart - Atrial Appendage | 18 | 290 | 5.8 |
| Adrenal Gland | 11 | 194 | 5.4 |
| Brain - Substantia nigra | 5 | 90 | 5.3 |
| Esophagus - Gastroesophageal Junction | 14 | 248 | 5.3 |
| Heart - Left Ventricle | 17 | 306 | 5.3 |
| Artery - Aorta | 16 | 295 | 5.1 |
| Brain - Amygdala | 5 | 102 | 4.7 |
| Brain - Cerebellar Hemisphere | 7 | 145 | 4.6 |
| Brain - Hypothalamus | 6 | 128 | 4.5 |
| Adipose - Subcutaneous | 16 | 454 | 3.4 |
| Muscle - Skeletal | 15 | 586 | 2.5 |

Yes=RPKM values above 0. No=RPKM values were 0. Per = percent of the samples with detectable expression. There were very few samples of some tissues, e.g., bladder.

**Supplemental Figure captions**

Supplemental Figure 1. RNA integrity (RIN) of samples of fungiform papillae (y-axis) versus nasal tissue (x-axis) for all 27 subjects. The blue line shows the linear correlation; grey shading shows the confidence interval around the correlation line.

**Supplemental Figure 2.** RNA-seq expressed as RPKM vs the cycle threshold from qPCR. Each dot is data from one person.

**Supplemental Figure 3.** Correlation among abundance of the *TAS2R38* PAV mRNA in nasal, taste tissue, and PTC taste intensity for all 19 heterozygous people providing samples. Each dot represents b subject (subject ID prefix “FUN”). The blue line shows linear correlation; the gray shading shows the confidence interval. Axes scales are the log of the relative abundance of the TAS2R8 PAV mRNA relative to *GAPDH*. Abundance of the *TAS2R38* PAV mRNA in nasal versus taste tissue (B) and PTC taste intensity ratings vs abundance of the *TAS2R38* PAV mRNA in taste tissue (E) and nasal tissue (C).

**Supplemental Figure 4.** PTC taste intensity ratings of subjects grouped by the *TAS2R38* diplotype (PAV is the active and AVI the inactive form). Each circle represents data from one person; red diamonds are the diplotype group mean.

**Supplemental Figure 5.** A violin plot of the *TAS2R38* mRNA expression by tissue from the GTEx dataset, dots represent the median value. Here we included data from all subjects and tissues including those with no *TAS2R38* expression (RPKM value = 0). The *TAS2R38* mRNA relative high expressed tissues are small intestine (terminal ileum), testis, pancreas, and colon (transverse).

**Supplemental Figure 6.** No significant difference (Kruskal-Wallis test, p > 0.05) in the *TAS2R38* mRNA expression among subjects with inactive form (AVI/AVI), active form (PAV/PAV) and heterozygous diplotype (AVI/PAV).

**Supplemental Figure 7.** Scatterplots of the *TAS2R38* mRNA expression across the tissues transverse colon, pancreas, small intestine (terminal ileum) and testis.

**Supplemental Figure 8.** Heatmap of the Spearman correlations of the *TAS2R38* mRNA expression across the tissues, colon (transverse), pancreas, small intestine (terminal ileum) and testis.

## References

1. Adler, E., et al., A novel family of mammalian taste receptors. Cell, 2000. 100: p. 693–702.

2. Chandrashekar, J., et al., T2Rs function as bitter taste receptors. Cell, 2000. 100(6): p. 703–11.

3. Matsunami, H., J.P. Montmayeur, and L.B. Buck, A family of candidate taste receptors in human and mouse. Nature, 2000. 404(6778): p. 601–4.

4. Lu, P., et al., Extraoral bitter taste receptors in health and disease. J Gen Physiol, 2017. 149(2): p. 181–197.

5. Lee, R.J., et al., T2R38 taste receptor polymorphisms underlie susceptibility to upper respiratory infection. J Clin Invest, 2012. 122(11): p. 4145–59.

6. Wooding, S., Phenylthiocarbamide: a 75-year adventure in genetics and natural selection. Genetics, 2006. 172(4): p. 2015–23.

7. Kim, U., et al., Worldwide haplotype diversity and coding sequence variation at human bitter taste receptor loci. Hum Mutat, 2005. 26(3): p. 199–204.

8. Risso, D.S., et al., Global diversity in the *TAS2R38* bitter taste receptor: revisiting a classic evolutionary PROPosal. Sci Rep, 2016. 6: p. 25506.

9. Bufe, B., et al., The molecular basis of individual differences in phenylthiocarbamide and propylthiouracil bitterness perception. Current Biology, 2005. 15(4): p. 322–7.

10. Lipchock, S.V., et al., Human bitter perception correlates with bitter receptor messenger RNA expression in taste cells. American Journal of Clinical Nutrition, 2013. 98: p. 1136–43.

11. Adappa, N.D., et al., Correlation of T2R38 taste phenotype and in vitro biofilm formation from nonpolypoid chronic rhinosinusitis patients. Int Forum Allergy Rhinol, 2016. 6(8): p. 783–91.

12. Mele, M., et al., The human transcriptome across tissues and individuals. Science, 2015. 348(6235): p. 660–5.

13. Brown, D.W., Smoking prevalence among US veterans. J Gen Intern Med, 2010. 25(2): p. 147–9.

14. Spielman, A.I., et al., Technique to collect fungiform (taste) papillae from human tongue. J Vis Exp, 2010. 18(42): p. 2201.

15. Lion, T., Appropriate controls for RT-PCR. Leukemia, 1996. 10(11): p. 1843.

16. Bolger, A.M., M. Lohse, and B. Usadel, Trimmomatic: a flexible trimmer for Illumina sequence data. Bioinformatics, 2014. 30(15): p. 2114–20.

17. Fu J, F.A., Collado-Torres L, Jaffe AE and Leek JT, ballgown: Flexible, isoform-level differential expression analysis. R package version 2.10.0, 2017.

18. Nuessle, T.M., et al., Denver papillae protocol for objective analysis of fungiform papillae. J Vis Exp, 2015(100): p. e52860.

19. Suitor, C.J., J. Gardner, and W.C. Willett, A comparison of food frequency and diet recall methods in studies of nutrient intake of low-income pregnant women. J Am Diet Assoc, 1989. 89(12): p. 1786–94.

20. Lipchock, S.V., et al., Caffeine bitterness is related to daily caffeine intake and bitter receptor mRNA abundance in human taste tissue. Perception, 2017: p. 301006616686098.

21. Livak, K.J. and T.D. Schmittgen, Analysis of relative gene expression data using real-time quantitative PCR and the 2(-Delta Delta C(T)) Method. Methods, 2001. 25(4): p. 402–8.

22. Pew Charitable Trust. Philadelphia: The state of the city. A 2014 update. 2014; Available from: http://www.pewtrusts.org/en/research-and-analysis/reports/2014/04/05/philadelphia-the-state-of-the-city-a-2014-update.

23. O’Mahony, M., et al., Confusion in the use of the taste adjectives ’sour’ and ’bitter’. Chemical Senses and Flavour, 1979. 4(4): p. 301–318.

24. Adappa, N.D., et al., The bitter taste receptor T2R38 is an independent risk factor for chronic rhinosinusitis requiring sinus surgery. Int Forum Allergy Rhinol, 2013.

25. Adappa, N.D., et al., Genetics of the taste receptor T2R38 correlates with chronic rhinosinusitis necessitating surgical intervention. Int Forum Allergy Rhinol, 2013.

26. Lin, B.P., Caries experience in children with various genetic sensitivity levels to the bitter taste of 6-n-propylthiouracil (PROP): a pilot study. Pediatr Dent, 2003. 25(1): p. 37–42.

27. Pidamale, R., et al., Genetic sensitivity to bitter taste of 6-n Propylthiouracil: A useful diagnostic aid to detect early childhood caries in pre-school children. Indian Journal of Human Genetics, 2012. 18(1): p. 101–5.

28. Oter, B., et al., The relation between 6-n-Propylthiouracil sensitivity and caries activity in schoolchildren. Caries Res, 2011. 45(6): p. 556–560.

29. Rupesh, S. and U.A. Nayak, Genetic sensitivity to the bitter taste of 6-n propylthiouracil: a new risk determinant for dental caries in children. J Indian Soc Pedod Prev Dent, 2006. 24(2): p. 63–8.

30. Chung, C.S., C.J. Witkop, and J.L. Henry, A genetic study of dental caries with special reference to PTC taste sensitivity. Am J Hum Genet, 1964. 16: p. 231–45.

31. The GTEx Consortium, The Genotype-Tissue Expression (GTEx) pilot analysis: multitissue gene regulation in humans. Science, 2015. 348(6235): p. 648–60.

32. Tomazic, P.V., et al., Nasal mucus proteomic changes reflect altered immune responses and epithelial permeability in patients with allergic rhinitis. J Allergy Clin Immunol, 2014. 133(3): p. 741–50.

33. Knowles, M.R. and R.C. Boucher, Mucus clearance as a primary innate defense mechanism for mammalian airways. J Clin Invest, 2002. 109(5): p. 571–7.

34. Behrens, M., et al., Immunohistochemical detection of *TAS2R38* protein in human taste cells. PLoS ONE, 2012. 7(7): p. e40304.

35. Jeon, T.I., et al., SREBP-2 regulates gut peptide secretion through intestinal bitter taste receptor signaling in mice. J Clin Invest, 2008. 118(11): p. 3693–700.

36. The Human Protein Atlas. [cited 2018 March 5th 2018]; Available from: https://www.proteinatlas.org/ENSG00000257138-TAS2R38/tissue/pancreas.

37. Delwiche, J.F., Z. Buletic, and P.A. Breslin, Relationship of papillae number to bitter intensity of quinine and PROP within and between individuals. Physiol Behav, 2001. 74(3): p. 329–37.

38. Bartoshuk, L.M., V.B. Duffy, and I.J. Miller, PTC/PROP tasting: Anatomy, psychophysics, and sex effects. Physiology and Behavior, 1994. 56: p. 1165–1171.

39. Kim, U.K., et al., Positional cloning of the human quantitative trait locus underlying taste sensitivity to phenylthiocarbamide. Science, 2003. 299(5610): p. 1221–5.

40. Timpson, N.J., et al., Refining associations between *TAS2R38* diplotypes and the 6-n-propylthiouracil (PROP) taste test: findings from the Avon Longitudinal Study of Parents and Children. BMC Genet, 2007. 8(1): p. 51.

41. Duffy, V.B., et al., Bitter receptor gene (*TAS2R38*), 6-n-propylthiouracil (PROP) bitterness and alcohol intake. Alcohol Clin Exp Res, 2004. 28(11): p. 1629–37.

42. Newcomb, R.D., M.B. Xia, and D.R. Reed, Heritable differences in chemosensory ability among humans. Flavour, 2012. 1(1): p. 9.

